# Evaluation of SARS-CoV-2 neutralizing antibodies using a vesicular stomatitis virus possessing SARS-CoV-2 spike protein

**DOI:** 10.1101/2020.08.21.262295

**Authors:** Hideki Tani, Long Tan, Miyuki Kimura, Yoshihiro Yoshida, Hiroshi Yamada, Shuetsu Fukushi, Masayuki Saijo, Hitoshi Kawasuji, Akitoshi Ueno, Yuki Miyajima, Yasutaka Fukui, Ippei Sakamaki, Yoshihiro Yamamoto, Yoshitomo Morinaga

## Abstract

SARS-CoV-2 is a novel coronavirus that emerged in 2019 and is now classified in the genus Coronavirus with closely related SARS-CoV. SARS-CoV-2 is highly pathogenic in humans and is classified as a biosafety level (BSL)-3 pathogen, which makes manipulating it relatively difficult due to its infectious nature. To circumvent the need for BSL-3 laboratories, an alternative assay was developed that avoids live virus and instead uses a recombinant VSV expressing luciferase and possesses the full length or truncated spike proteins of SARS-CoV-2. Furthermore, to measure SARS-CoV-2 neutralizing antibodies under BSL2 conditions, a chemiluminescence reduction neutralization test (CRNT) for SARS-CoV-2 was developed. The neutralization values of the serum samples collected from hospitalized patients with COVID-19 or SARS-CoV-2 PCR-negative donors against the pseudotyped virus infection evaluated by the CRNT were compared with antibody titers determined from an immunofluorescence assay (IFA). The CRNT, which used whole blood collected from hospitalized patients with COVID-19, was also examined. As a result, the inhibition of pseudotyped virus infection was specifically observed in both serum and whole blood and was also correlated with the results of the IFA. In conclusion, the CRNT for COVID-19 is a convenient assay system that can be performed in a BSL-2 laboratory with high specificity and sensitivity for evaluating the occurrence of neutralizing antibodies against SARS-CoV-2.

## Introduction

Recently, the infectious Coronavirus Disease 2019 (COVID-19) emerged and is caused by a newly identified coronavirus, severe acute respiratory syndrome coronavirus 2 (SARS-CoV-2) (1). Globally, COVID-19 severely impacted health and socio-economic conditions. Currently, no safe and effective antivirals or other therapies for COVID-19 exist, although some drugs, such as remdesivir, showed limited efficacy for the treatment of patients with COVID-19 (2, 3). Similar to other diseases, proper treatment requires an accurate diagnosis. Therefore, establishing diagnostics, such as the detection of target viral genes and antibodies, is also required.

The genome structure of SARS-CoV-2 is similar to that of severe acute respiratory syndrome coronavirus SARS-CoV, which is the causative agent for severe acute respiratory syndrome (SARS) that showed high mortality and morbidity in the late 2002-to the middle of 2003 outbreak, which mainly occurred in China (4). The proteins of SARS-CoV-2 consist of two large polyproteins: ORF1a and ORF1ab; four structural proteins: spike (S), envelope (E), membrane (M), and nucleocapsid (N); and eight accessory proteins: ORF3a, ORF3b, ORF6, ORF7b, ORF8a, ORF8b, and ORF9b. S protein is a glycoprotein, which is responsible for binding and penetration of target cells. The S protein is also important for induction of protective humoral and cellular immunity during infection. The S protein is the main target with which the neutralizing antibodies react. Measuring the SARS-CoV-2 neutralizing antibodies is important for proper diagnosis, to study the serological epidemiology and determine infection control of SARS-CoV-2.

The enzyme-linked immunosorbent assay (ELISA), immunofluorescence assay (IFA), and immunochromatography utilize the principle of antigen-antibody reaction and were developed as serodiagnostic methods for SARS-CoV-2 infection.

The specificity of these assays is fairly high, but problems are present such as relatively low sensitivity and a high rate of false positives. The neutralization antibody test (NT) for serum using live SARS-CoV-2 is a method in which inhibition of the serum upon viral infection is observed in the presence of neutralizing antibodies against proteins involved in viral binding and penetration in the serum. Generally, the NT is the standard method used to confirm the presence of neutralizing antibodies against SARS-CoV-2. However, this method takes a long time because it depends on the growth of the virus, and it is not a simple measuring system in terms of complicated handling and biosafety for highly pathogenic SARS-CoV-2. Therefore, recently, a pseudotyped virus system based on vesicular stomatitis virus (VSV) or pseudotyped particle systems based on lentivirus or retrovirus were developed for the detection of neutralizing antibodies instead of using infectious and authentic viruses (5–7). However, a significant need exists to evaluate the neutralizing antibody against SARS-CoV-2 in the clinical setting. Therefore, the ability to perform the NT under lower BSL laboratory conditions is preferred.

In this study, an antibody detection system based on the chemiluminescence reduction neutralization test (CRNT) and using the S protein-based pseudotyped viruses was developed. The correlation between antibody titers against SARS-CoV-2 determined by CRNT were evaluated along with those determined by the IFA in which recombinant S protein was used as an antigen. The usefulness of the CRNT system for detecting neutralizing antibodies against SARS-CoV-2 that was newly developed in this study is presented.

## Materials and Methods

### Plasmids

The cDNAs of the SARS-CoV-2 spike protein were obtained by chemical synthesis with optimization for the humanized codon (Integrated DNA Technologies, Inc., Coralville, IA). The S cDNA of SARS-CoV-2 was cloned into the pCAGGS expression vector (8). The resulting plasmid was designated as pCAG-SARS-CoV-2. The plasmid, which contains the S protein gene with a 19 aa truncation at the C-terminus, was constructed using the cDNA of pCAG-SARS-CoV-2. The S proteins with the 19 aa deletion of coronaviruses were previously reported to show increased efficiency regarding incorporation into virions of VSV (9, 10).

### Cells

Human (Huh7 and 293T), monkey (Vero), hamster (BHK and CHO), and mouse (NIH3T3) cell lines were obtained from the American Type Culture Collection (Summit Pharmaceuticals International, Tokyo, Japan). All cell lines were grown in Dulbecco’s modified Eagle’s medium (DMEM; Nacalai Tesque, Inc., Kyoto, Japan) containing 10% heat inactivated fetal bovine serum (FBS).

### Generation of pseudotyped VSVs

Pseudotyped VSVs bearing the S protein, the 19 aa-truncated S protein of SARS-CoV-2, or VSV-G were generated as described below. Briefly, 293T cells were grown to 80% confluence on collagen-coated tissue culture plates and then transfected with each expression vector: pCAG-SARS-CoV-2 S-full, pCAG-SARS-CoV-2 S-t19, and pCAG-VSV-G. After 24 h of incubation, the cells transfected with each plasmid were infected with G-complemented (*G) VSVΔG/Luc (*G-VSVΔG/Luc)(11) at a multiplicity of infection (MOI) of 0.5 per cell. Then, the virus was adsorbed and extensively washed four times with 10% FBS DMEM. After 24 h of incubation, to remove cell debris, the culture supernatants containing pseudotyped VSVs were centrifuged, and then, they stored at −80°C until ready for use. The pseudotyped VSV bearing SARS-CoV-2 S protein or SARS-CoV-2 truncated S protein are referred to as Sfullpv or St19pv, respectively. The infectivity of Sfullpv, St19pv, or VSVpv to 293T cells was assessed by measuring the luciferase activity. The value of the relative light unit (RLU) of luciferase was determined using a PicaGene Luminescence Kit (TOYO B-Net Co., LTD, Tokyo, Japan) and GloMax Navigator System G2000 (Promega Corporation, Madison, WI), according to the manufacturer’s protocol.

### Coomassie Brilliant Blue (CBB) staining and Immunoblotting

Transfection of 293T cells occurred with pCAG-SARS-CoV-2 Sfull, pCAG-SARS-CoV-2 St19, or VSV-G. At 24 h post-transfection, the cells were collected and lysed in phosphate-buffered saline (PBS) containing 1% NP40. Then, the lysates were centrifuged to separate insoluble pellets from supernatants. The supernatants were used as samples. The Sfullpv or St19pv, which were generated as described above, were pelleted through a 20% (wt/vol) sucrose cushion at 25,000 rpm for 2 h in an SW41 rotor (Beckman Coulter, Tokyo, Japan). Then, the pellets were resuspended in PBS. Each sample that was boiled in loading buffer was subjected to 10% sodium dodecyl sulfate-polyacrylamide gel electrophoresis (SDS-PAGE). According to the manufacturer’s protocol, the proteins in the gel were stained with CBB Stain One (Nacalai Tesque, Inc.). Next, the proteins in another gel were electrophoretically transferred to a methanol-activated polyvinylidene difluoride (PVDF) membrane (Millipore, Billerica, MA) and reacted with COVID-19 hospitalized patient sera (#12). Then, immune complexes were visualized with SuperSignal West Dura Extended Duration Substrate (Pierce, Rockford, IL) and detected by an LAS3000 analyzer (Fuji Film, Tokyo, Japan).

### Blood samples

Twenty-three serum samples were collected from hospitalized patients with COVID-19 who were admitted to the University of Toyama Hospital, Toyama, Japan. In addition, nineteen serum samples were collected from COVID-19 PCR-negative donors at the University of Toyama Hospital. The diagnosis of COVID-19 in all patients or donors was assessed using the real-time PCR method with specific primers, which were developed at the National Institute of Infectious Diseases, Japan (12).

By using a blood collection tube containing EDTA, whole blood samples were obtained from 5 hospitalized patients (the University of Toyama Hospital) with COVID-19.

### Neutralization assays with patient blood samples

The patient sera used in this study were collected from participants after obtaining informed consent. To examine neutralization of the human serum or whole blood samples against pseudotyped viruses, Vero cells were treated with serially diluted sera or whole blood of convalescent patients with COVID-19 or PCR-negative donors and then inoculated with Sfullpv, St19pv, or VSVpv. To remove hematopoietic cells from whole blood samples, centrifugation was performed at 2,000 × g for 5 min. Infectivity of the pseudotyped viruses were determined by measuring luciferase activities after 24 h of incubation at 37 °C.

### Immunofluorescence assay (IFA)

For the IFA, BHK cells transfected with pCAG-SARS-CoV-2 S-full were fixed with acetone-methanol (1:1) at 4 °C for 20 min. Fixed cells were reacted with the test serum samples, which were diluted at 1:100 with PBS. After an incubation for 1 h, the cells were rinsed with PBS and incubated with goat anti-human Alexa Fluor 488 (Invitrogen). After washing with PBS, staining was observed under a fluorescence microscope.

### Ethics statement

All of the samples, protocols, and procedures were approved by the Research Ethics Committee at the University of Toyama for the use of human subjects (approval number: R2019167).

## Results

### Production and characterization of SARS-CoV-2 Sfullpv and St19pv

SARS-CoV-2 Sfullpv, St19pv, and VSVpv were generated in 293T cells transiently expressed with full length-, truncated S proteins of SARS-CoV-2 and VSV-G, respectively, upon infection of 293T cells with *G-VSVΔG/Luc, as previously reported (11). To confirm the incorporation of S proteins into Sfullpv or St19pv particles, the pseudotyped viruses were purified by ultracentrifugation and analyzed using immunoblotting and the serum of COVID-19 hospitalized patient (Fig. 1A, right panel). The S1 and S2 proteins in Sfullpv or St19pv and Sfull- or St19-expressing cell lysates were detected but not in VSV that lacked an envelope protein (ΔGpv) or mock cells (Ctrl). The VSV structural proteins, nucleoprotein (N), and matrix protein (M) were also detected in Sfullpv, St19pv, and ΔGpv by CBB staining (Fig. 1A, left panel). Overall, the amount of St19 proteins incorporated was higher than that of Sfull proteins, although the amount of structural proteins of VSV was almost the same level among all the virions. These results indicate that the incorporation of truncated S proteins into VSV particles was more efficient than the full-length S protein. Next, ΔGpv, Sfullpv, St19pv, and VSVpv were inoculated into the indicated cell lines to examine the infectivity of pseudotyped viruses to various mammalian cell lines (Fig. 1B). Among the tested cell lines, Huh7, 293T, Vero, and BHK cells were susceptible to Sfullpv and St19pv infection. NIH3T3 cells were less susceptible to infection by both Sfullpv and St19pv, while CHO cells showed no susceptibility. Notably, the infectivity of St19pv was higher than that of Sfullpv in Vero cells.

**Fig. 1.**
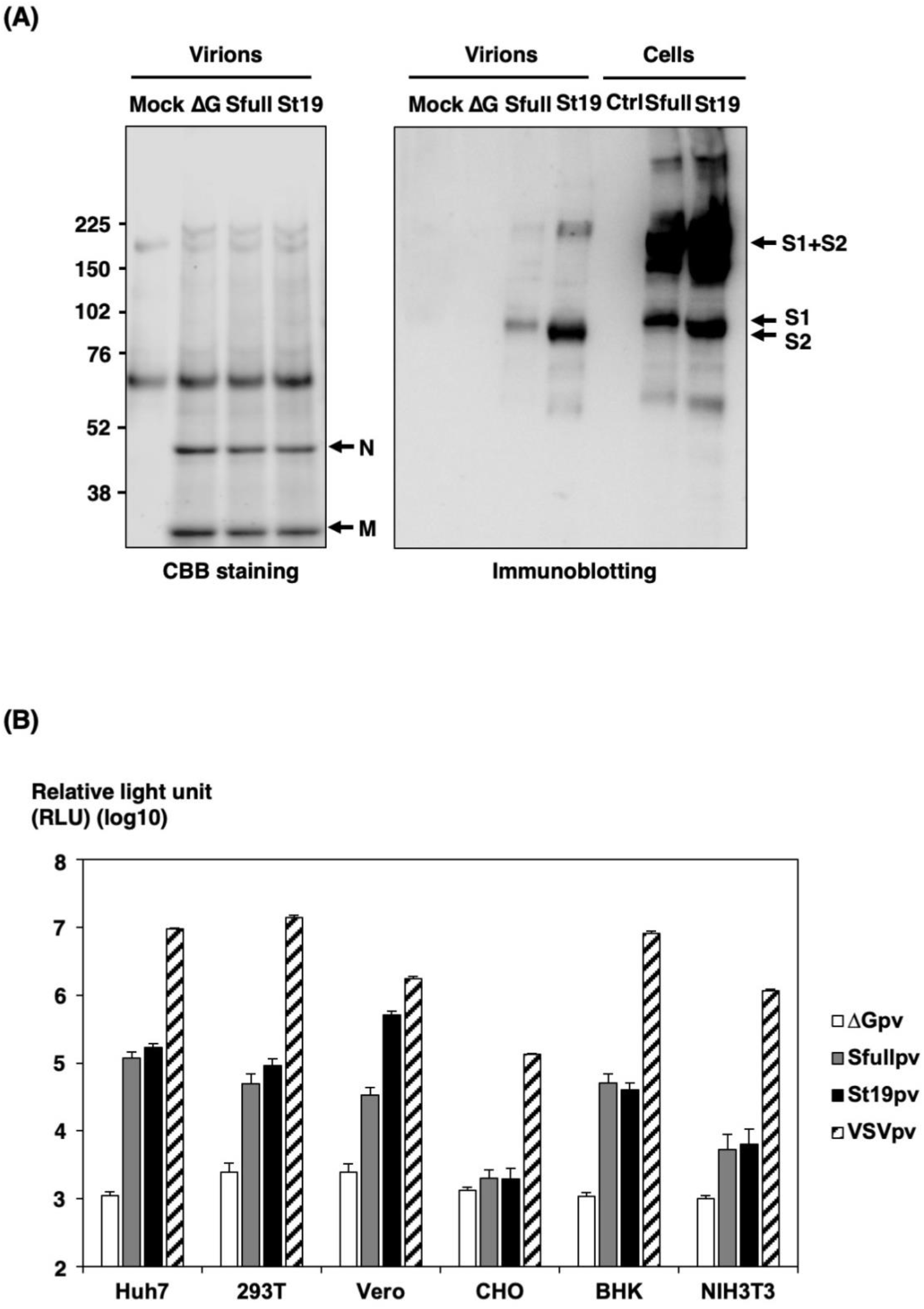
Characterization of pseudotyped viruses possessing spike proteins of SARS-CoV-2. (A) Incorporation or expression of S proteins in the virions or cells were investigated using CBB staining and immunoblotting. (B) Efficiency of gene transduction in various mammalian cell lines using the pseudotyped viruses. Sfullpv, St19pv, and 100-fold-diluted VSVpv generated in 293T cells were inoculated into the indicated cell lines. At 24 h post-infection, infectivity of the viruses was determined by measuring luciferase activities as a relative luciferase unit (RLU). VSVpv without envelope (ΔGpv) was used as a negative control. The results are from three independent assays with error bars representing standard deviations.

To determine the specificity of infection of Sfullpv and St19pv, a neutralization assay of the pseudotyped viruses was performed using sera of two hospitalized COVID-19 patients. The infectivity of Sfullpv and St19pv, but not that of VSVpv in Vero cells, were clearly inhibited by sera of the patients in a dose-dependent manner (Fig. 2). These data indicated that Sfullpv and St19pv infection exhibited an S protein-mediated entry.

**Fig. 2.**
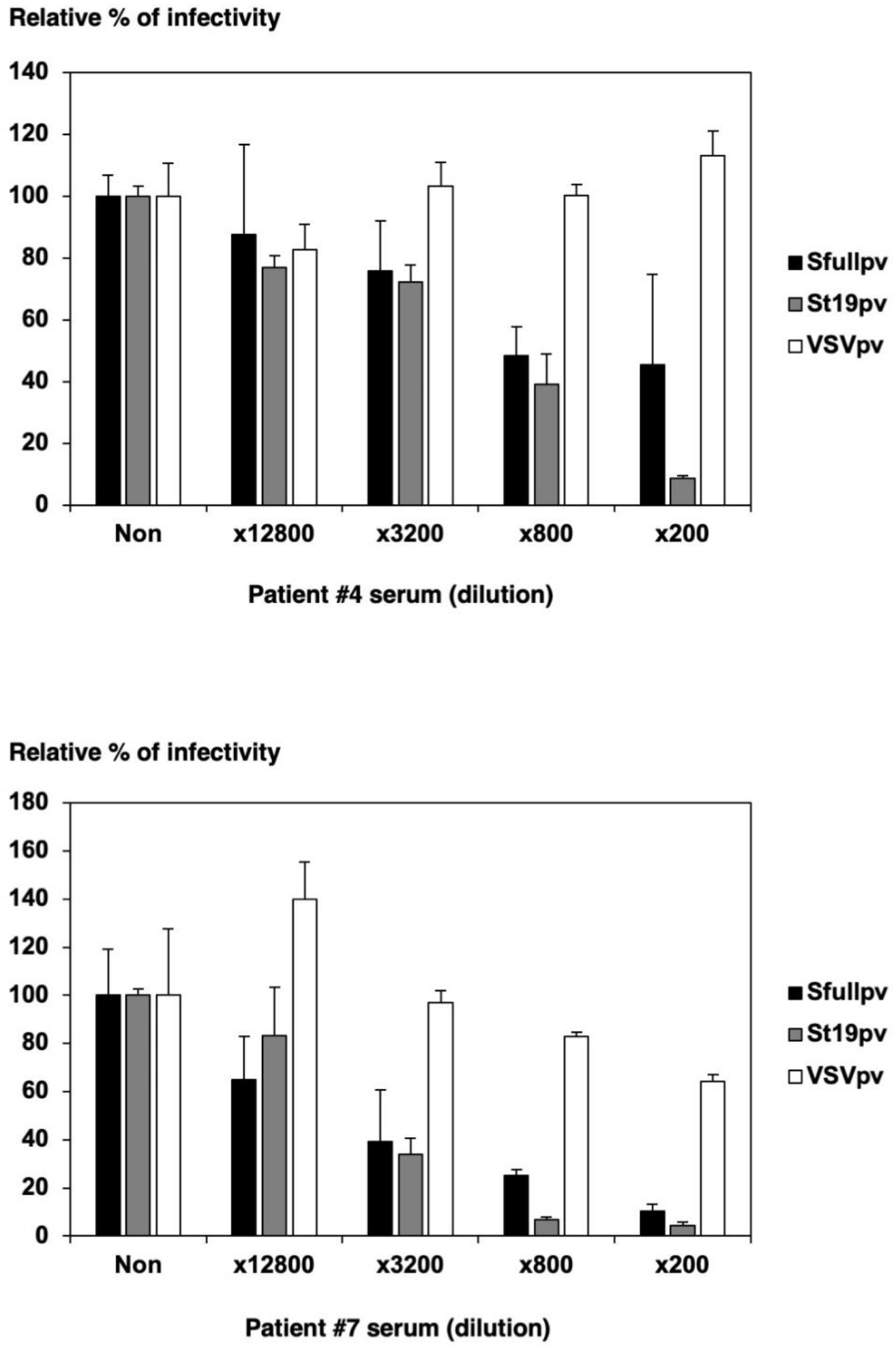
Dose-dependent effects on the neutralization of Sfullpv and St19pv infections through sera from convalescent COVID-19 patients. The pseudotyped viruses were preincubated with the indicated dilutions of sera from two different convalescent COVID-19 patients (patients #4 and #7). Thereafter, Vero cells were infected with pseudotyped viruses. Infectivities of pseudotyped viruses were determined by measuring luciferase activities at 24 h post-infection. The results are from three independent assays with error bars representing standard deviations.

### Neutralization test for COVID-19 hospitalized patients or COVID-19 PCR-negative donors by pseudotyped viruses

To examine the neutralization of COVID-19 hospitalized patients or COVID-19 PCR-negative donors against St19pv, Vero cells with each serum were infected with St19pv and VSVpv.

Neutralization of St19pv was observed based on the sixteen sera of COVID-19 hospitalized patients at a rate of more than 99% (Fig. 3A). Sera, which did not show the neutralization of St19pv, were derived from COVID-19 hospitalized patients who were hospitalized for a short period before antibody production (such as within 3 days after onset). No neutralization was observed in the VSVpv infection by any of the sera of COVID-19 hospitalized patients and in both St19pv and VSVpv infection by any of the sera of COVID-19 PCR-negative donors (Fig. 3A and B). The dot plot graph shows a classification of each pseudotyped virus from Fig. 3A and 3B graphs (Fig. 3C). Due to the presence of convalescent and non-convalescent patient sera, the degree of neutralization activity of St19pv by COVID-19 hospitalized patient sera was variable.

**Fig. 3.**
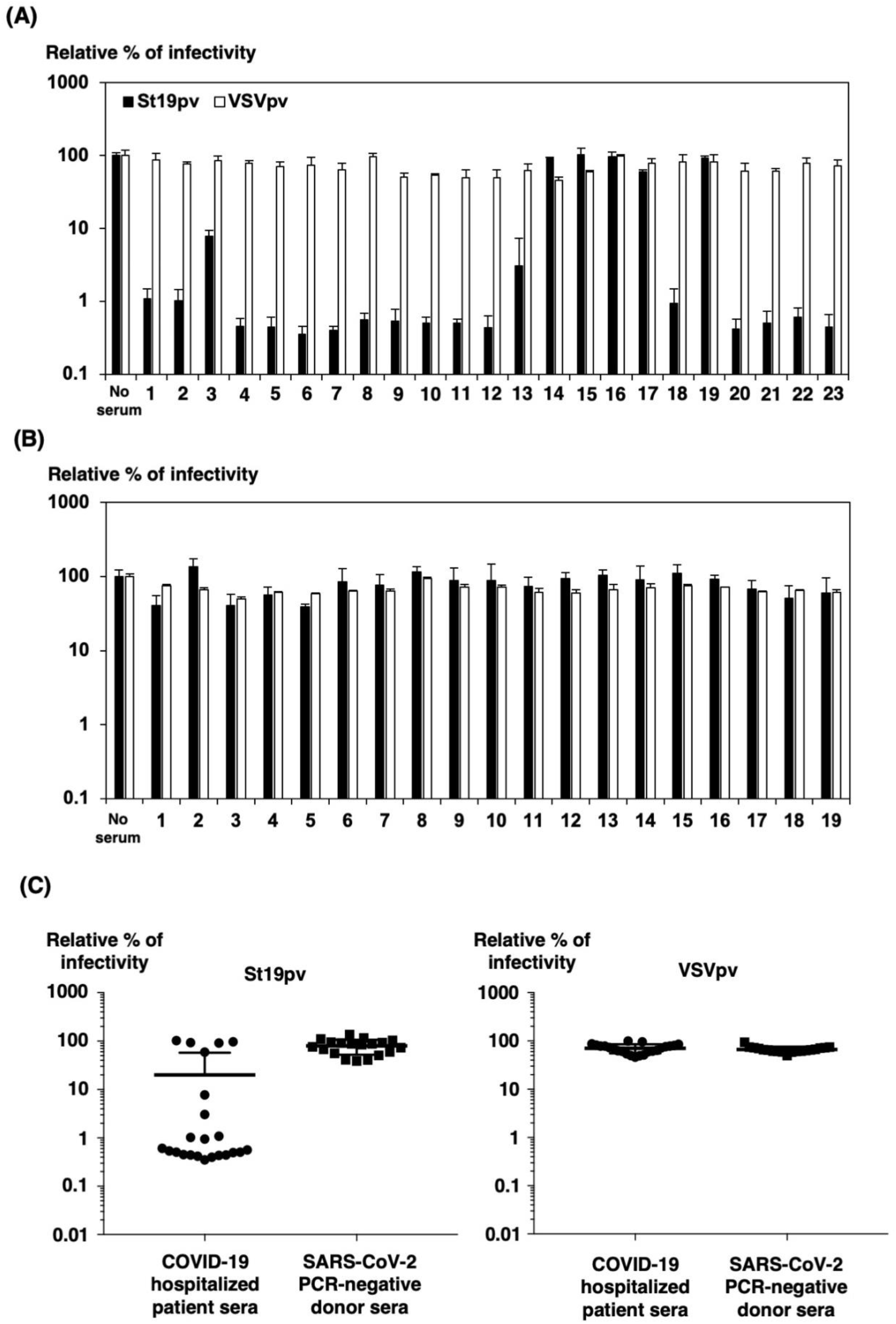
Neutralization of St19pv and VSVpv infections through sera from hospitalized patients with COVID-19 or a SARS-CoV-2 PCR-negative donor. The pseudotyped viruses were preincubated with two a hundred-fold dilution of sera from 23 hospitalized COVID-19 patients (A) or 19 SARS-CoV-2 PCR-negative donors (B). Thereafter, Vero cells were infected with pseudotyped viruses. Infectivities of pseudotyped viruses were determined by measuring luciferase activities 24 h post-infection. The results are from three independent assays with error bars representing standard deviations. (C) A summary of the results of (A) and (B) divided by pseudotyped viruses are represented by dot plot analyses.

### IFA for COVID-19 hospitalized patients or COVID-19 PCR-negative donors

To examine the correlation of the antibody titers using CRNT compared to those determined by the IFA, the IFA was also performed using COVID-19 hospitalized patients or COVID-19 PCR-negative donors. The fluorescence intensity of IFA was correlated with the sera, which exhibit a high neutralizing activity in the CRNT (Fig. 3 and Table 1). Sera with low neutralizing activity in the CRNT showed weak fluorescence intensity, and sera that demonstrated no neutralizing activity in the CRNT were also negative by IFA.

**Table 1.** Intensity of IF positive cells expressing SARS-CoV-2 S protein.

| Hospitalized COVID-19 patient sera |  | SARS-CoV-2-negative donor sera |  |
| --- | --- | --- | --- |
| No. | IF-Intensity | No. | IF-Intensity |
| 1 | +++ | 1 | – |
| 2 | +++ | 2 | – |
| 3 | ++ | 3 | – |
| 4 | +++ | 4 | – |
| 5 | +++ | 5 | – |
| 6 | +++ | 6 | – |
| 7 | +++ | 7 | – |
| 8 | +++ | 8 | – |
| 9 | +++ | 9 | – |
| 10 | ++ | 10 | – |
| 11 | +++ | 11 | – |
| 12 | +++ | 12 | – |
| 13 | – | 13 | – |
| 14 | – | 14 | – |
| 15 | – | 15 | – |
| 16 | – | 16 | – |
| 17 | – | 17 | – |
| 18 | + | 18 | – |
| 19 | – | 19 | – |
| 20 | + |  |  |
| 21 | ++ |  |  |
| 22 | ++ |  |  |
| 23 | ++ |  |  |

### Comparison of patient sera and whole blood in the neutralization of pseudotyped viruses

Also, we compared the neutralizing effect of pseudotyped viruses between the sera and whole blood of COVID-19 hospitalized patients with or without centrifugation (Fig. 4). After centrifugation of whole blood, hematopoietic cells, including red blood cells, may be removed. As a result, the neutralization of both sera and whole blood against St19pv infection was observed with or without centrifugation (Fig. 4). Although the neutralizing activities of whole blood were higher than those of the sera in St19pv infection, the infectivity of VSVpv was reduced by approximately 1/10 without centrifugation (Fig. 4A). The reduction of infectivity of both St19pv and VSVpv was suppressed, by removing hematopoietic cells in whole blood after centrifugation (Fig. 4B). Therefore, some inhibitory factors, such as hematopoietic cells, may be involved in pseudotyped virus infection in whole blood.

**Fig. 4.**
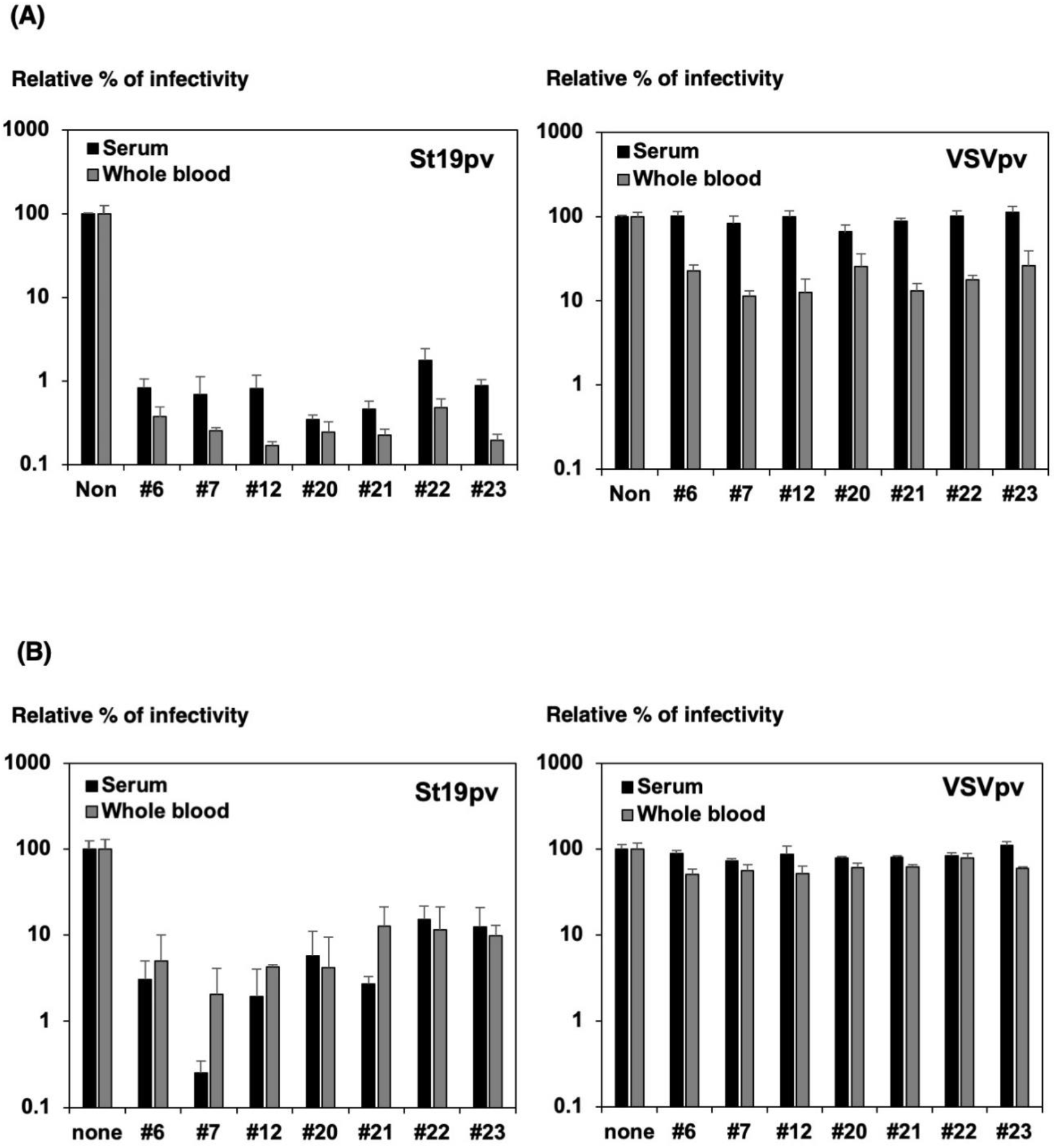
Effects on the neutralization of St19pv infection by serum or whole blood from COVID-19 hospitalized patients. The pseudotyped viruses were preincubated with two hundred dilutions of sera from seven different COVID-19 hospitalized patients (patients #6, #7, #12, #20, #21, #22, and #23). Thereafter, Vero cells were infected with pseudotyped viruses. (A) Infectivities of pseudotyped viruses treated with serum or whole blood without centrifugation. (B) Infectivities of pseudotyped viruses treated with serum or whole blood after centrifugation. Infectivities of pseudotyped viruses were determined by measuring luciferase activities 24 h post-infection. The results are from three independent assays with error bars representing standard deviations.

## Discussion

A rapid, safe, and highly sensitive CRNT system using VSV-based pseudotyped viruses with SARS-CoV-2 S or truncated S proteins was developed. Because this system utilizes replication and translation of VSV, neutralization against pseudotyped virus infection can be determined within 12–16 hours. Another pseudotyped viral system that uses retroviral or lentiviral vectors takes approximately 48 h to obtain results. Therefore, the VSV-based pseudotyped viral system is considered more useful. In addition, since measurement of luciferase activity is a quantitative method, it is not necessary to count GFP-positive cells. Therefore, this CRNT system permits a simple and objective evaluation for the neutralization.

For many viral species, CRNT systems were developed using VSV-pseudotyped viruses with their own envelope proteins (11, 13–15). In SARS-CoV-2, researchers recently demonstrated the construction of pseudotyped viruses and evaluation of the presence of neutralizing antibodies (16).

In this study, we prepared a pseudotyped virus that possesses a truncated SARS-CoV-2 S protein, which showed higher infectivity. Furthermore, the neutralizing activity of the test sera and whole blood against the pseudotyped virus was quantitatively detected in a convalescent patient with COVID-19, while the donor sera of the COVID-19 PCR-negative patient showed a negative reaction. In the CRNT of St19pv, the infectivity of St19pv was reduced by 99% or more by the convalescent phase patient sera. This demonstrated that the convalescent phase patient sera of COVID-19 exhibited a high neutralizing antibody activity. The results determined by the CRNT also correlated with those determined by the IFA. Antibodies against the S protein of SARS-CoV-2 in COVID-19 convalescent patient sera were capable of neutralizing the viral infection.

If using whole blood in the CRNT becomes possible, the work of separating serum will no longer be necessary. The CRNT can be performed with an extremely small amount of blood sample (only a few microliters). Therefore, when we confirmed that the CRNT with whole blood of the convalescent phase patient of COVID-19 was possible, a high neutralizing activity by the CRNT should be observed. However, the infectivity of the control VSVpv was also reduced by the whole blood control. Since many hematopoietic cells, including red blood cells, are contained in whole blood, these cells are present on the Vero cells used in the CRNT with whole blood. The possibility highly exists that this inhibition is due to the presence of hematopoietic cells because removal of the cells with centrifugation suppressed non-specific reduction of the pseudotyped viral infection.

However, inhibition of St19pv infection shows a stronger neutralizing activity compared to VSVpv infection. Therefore, evaluating the CRNT using whole blood is possible.

The neutralizing antibody measurement system using pseudotyped viruses for SARS-CoV-2 is an effective tool for evaluating the presence or duration of the neutralizing antibody in convalescent patients and to screen for those who present with the neutralizing antibody among suspected populations. In addition, this CRNT system does not require the use of infectious viruses to measure neutralizing antibodies. Therefore, once the pseudotyped virus system is established, it can be made available at many laboratories without BSL-3 facilities. Furthermore, because of the measuring system by chemiluminescence, the results can be obtained safely and quickly. Finally, the CRNT using whole blood is a simpler and safer method because it can be measured with only a very small amount of blood from an eligible person.

## Acknowledgments

We gratefully acknowledge Ms. Kaoru Hounoki and Yoriko Ito for technical and secretarial assistance. This work was supported in part by a grant-in-aid from the Japan Society for the Promotion of Science (JSPS KAKENHI Grant Number JP18K07144), and by a grant-in-aid from the Japan Agency for Medical Research and Development (AMED under Grant Number JP20fk0108081), and a grant from the Takeda Science Foundation.

## References

1. Huang C, Wang Y, Li X, Ren L, Zhao J, Hu Y, Zhang L, Fan G, Xu J, Gu X, Cheng Z, Yu T, Xia J, Wei Y, Wu W, Xie X, Yin W, Li H, Liu M, Xiao Y, Gao H, Guo L, Xie J, Wang G, Jiang R, Gao Z, Jin Q, Wang J, Cao B. 2020. Clinical features of patients infected with 2019 novel coronavirus in Wuhan, China. Lancet 395:497–506.

2. Musa A, Pendi K, Hashemi A, Warbasse E, Kouyoumjian S, Yousif J, Blodget E, Stevens S, Aly B, Baron DA. 2020. Remdesivir for the treatment of COVID-19: A systematic review of the literature. West J Emerg Med 21:737–741.

3. Tu H, Tu S, Gao S, Shao A, Sheng J. 2020. Current epidemiological and clinical features of COVID-19; a global perspective from China. J Infect 81:1–9.

4. Jin Y, Yang H, Ji W, Wu W, Chen S, Zhang W, Duan G. 2020. Virology, epidemiology, pathogenesis, and control of COVID-19. Viruses 12:372.

5. Li Q, Liu Q, Huang W, Li X, Wang Y. 2018. Current status on the development of pseudoviruses for enveloped viruses. Rev Med Virol 28.

6. Joglekar AV, Sandoval S. 2017. Pseudotyped lentiviral vectors: One vector, many Guises. Hum Gene Ther Methods 28:291–301.

7. Whitt MA. 2010. Generation of VSV pseudotypes using recombinant DeltaG-VSV for studies on virus entry, identification of entry inhibitors, and immune responses to vaccines. J Virol Methods 169:365–74.

8. Saijo M, Qing T, Niikura M, Maeda A, Ikegami T, Prehaud C, Kurane I, Morikawa S. 2002. Recombinant nucleoprotein-based enzyme-linked immunosorbent assay for detection of immunoglobulin G antibodies to Crimean-Congo hemorrhagic fever virus. J Clin Microbiol 40:1587–91.

9. Fukushi S, Mizutani T, Saijo M, Matsuyama S, Miyajima N, Taguchi F, Itamura S, Kurane I, Morikawa S. 2005. Vesicular stomatitis virus pseudotyped with severe acute respiratory syndrome coronavirus spike protein. J Gen Virol 86:2269–2274.

10. Kawase M, Shirato K, Matsuyama S, Taguchi F. 2009. Protease-mediated entry via the endosome of human coronavirus 229E. J Virol 83:712–21.

11. Tani H, Shiokawa M, Kaname Y, Kambara H, Mori Y, Abe T, Moriishi K, Matsuura Y. 2010. Involvement of ceramide in the propagation of Japanese encephalitis virus. J Virol 84:2798–807.

12. Shirato K, Nao N, Katano H, Takayama I, Saito S, Kato F, Katoh H, Sakata M, Nakatsu Y, Mori Y, Kageyama T, Matsuyama S, Takeda M. 2020. Development of genetic diagnostic methods for detection for novel coronavirus 2019(nCoV-2019) in Japan. Jpn J Infect Dis 73.

13. Sakata M, Tani H, Anraku M, Kataoka M, Nagata N, Seki F, Tahara M, Otsuki N, Okamoto K, Takeda M, Mori Y. 2017. Analysis of VSV pseudotype virus infection mediated by rubella virus envelope proteins. Sci Rep 7:11607.

14. Tani H, Iha K, Shimojima M, Fukushi S, Taniguchi S, Yoshikawa T, Kawaoka Y, Nakasone N, Ninomiya H, Saijo M, Morikawa S. 2014. Analysis of Lujo virus cell entry using pseudotype vesicular stomatitis virus. J Virol 88:7317–30.

15. Tani H, Shimojima M, Fukushi S, Yoshikawa T, Fukuma A, Taniguchi S, Morikawa S, Saijo M. 2016. Characterization of glycoprotein-mediated entry of severe fever with thrombocytopenia syndrome Virus. J Virol 90:5292–301.

16. Crawford KHD, Eguia R, Dingens AS, Loes AN, Malone KD, Wolf CR, Chu HY, Tortorici MA, Veesler D, Murphy M, Pettie D, King NP, Balazs AB, Bloom JD. 2020. Protocol and reagents for pseudotyping lentiviral particles with SARS-CoV-2 spike protein for neutralization assays. Viruses 12:513.

